# Xport-A functions as a chaperone by stabilizing the first 5 transmembrane domains of Rhodopsin-1

**DOI:** 10.1101/2022.04.01.486763

**Authors:** Catarina J Gaspar, Joana C Martins, Manuel N Melo, Tiago N Cordeiro, Colin Adrain, Pedro M Domingos

## Abstract

Rhodopsin-1 (Rh1), the main photo-sensitive protein of *Drosophila*, is a seven transmembrane domain protein, which is inserted co-translationally in the endoplasmic reticulum (ER) membrane. Maturation of Rh1 occurs in the ER, where various chaperones interact with Rh1 to aid in its folding and subsequent transport in the secretory pathway. Xport-A has been shown to be a chaperone/ transport factor for Rh1, but the exact molecular mechanism for Xport-A activity upon Rh1 is not known. Here, based on computational predictions, we propose a model where Xport-A functions as a chaperone in the biosynthesis of Rh1 by stabilizing the first 5 transmembrane domains of Rh1, but not the full length Rh1 protein.

## Introduction

Rh1 functional protein is composed by the apoprotein opsin and the covalently-bound chromophore 11-*cis* 3-hydroxy-retinal (Ozaki *et al*, 1993). The opsin corresponds to the protein moiety of Rh1 and is encoded by the ninaE (neither inactivation nor afterpotential E) gene (O’Tousa *et al*, 1985; Zuker *et al*, 1985). This gene encodes an integral membrane protein composed of seven TMDs (O’Tousa *et al*, 1985), which is inserted co-translationally into the ER membrane. Rh1 maturation in the ER involves post-translational modifications such as transient glycosylation, followed by de-glycosylation (O’Tousa, 1992; Katanosaka *et al*, 1998; Webel *et al*, 2000) and chromophore binding (Ozaki *et al*,1993). Then, Rh1 is transported through Golgi compartments to the rhabdomeres, where phototransduction takes place (Colley *et al*, 1991; Wolff & Ready, 1991).

Rh1 requires several chaperones for proper folding and transport out of the ER, including ninaA (neither inactivation nor afterpotential A), Calnexin (Cnx), Xport-A and Xport-B. The ninaA mutant was first identified by abnormal electroretinograms (ERGs) presenting “neither inactivation nor afterpotential” that is characteristic of mutants with reduced levels of functional rhodopsin (Pak *et al*,1970). NinaA is a homolog of the vertebrate cyclophilin, a target of the immunosuppressant drug cyclosporine that presents *cis-trans-isomerase* activity (Schneuwly *et al*, 1989), but does not seem to function as an enzyme in *Drosophila* eyes, and functions instead as a Rh1 chaperone (Colley *et al*, 1991). NinaA is predominantly localized to the ER and secretory vesicles, and forms a complex with Rh1 to promote maturation and transport of Rh1 through the secretory pathway (Colley *et al*, 1991; Baker *et al*, 1994). In ninaA mutants, there is a substantial reduction of Rh1 levels and an ER accumulation of immature glycosylated Rh1, resulting in ER expansion (Colley *et al*, 1991). These mutants do not display obvious photoreceptor degeneration (Rosenbaum *et al*, 2006), which could be explained by degradation of immature Rh1 through ERAD or other cellular mechanisms (Xiong & Bellen, 2013).

Cnx is an ER-resident type I membrane protein that binds glycosylated proteins in a Ca^2+^ dependent manner, to aid in protein folding (Pearse & Hebert, 2010). In *Drosophila* photoreceptor cells, Cnx physically interacts with Rh1 and is essential for its maturation (Rosenbaum *et al*, 2006). Loss of Cnx results in extensive reduction of functional Rh1 at the rhabdomeres, with a small fraction of Rh1 in an immature glycosylated state. Cnx mutations present an age-related retinal degeneration, accompanied by accumulations of ER cisternae and various types of deposits, consistent with a failure in Rh1 maturation. This phenotype was shown to be light-dependent and this dependency has been explained by the role that Cnx plays in Ca^2+^ buffering in the cell body. These results indicate that Cnx plays a dual role in maintaining photoreceptor cell survival, by promoting Rh1 maturation and regulating Ca^2+^ levels (Rosenbaum *et al*, 2006).

The Xport locus is bicistronic; it is transcribed as a single mRNA that encodes two different proteins: Xport-A and Xport-B (Chen *et al*, 2015). Xport-A is a tail-anchored (TA) protein and Xport-B is predicted to be a type III membrane protein. Both proteins have homologs in insect species (Rosenbaum *et al*, 2011; Chen *et al*, 2015), and the bicistronic nature of the locus is also conserved (Chen *et al*, 2015). Xport-A was the first of these proteins to be described (Rosenbaum *et al*, 2011), in a screening of the Zuker collection of EMS-mutagenized flies. Xport-A mutant (Xport-A^1^) homozygous flies presented an ERG profile similar to mutants of the TRP (transient receptor potential) channel and displayed extremely reduced levels of both Rh1 and TRP proteins. The mutation in Xport-A^1^ is recessive, as the heterozygotes presented normal protein levels of TRP and Rh1. When reared in the dark, Xport-A^1^ flies exhibited ER accumulations and Golgi expansion, but the rhabdomere morphology was preserved. Upon exposure to light, the defect worsened and the mutants presented retinal degeneration. These results indicate that the retinal degeneration in Xport-A mutants results from the combined effects of protein aggregation (due to ER retention of Rh1 and TRP), which cause light-independent effects, and misregulation of Ca^2+^ levels (due to loss of TRP), which causes the light-enhanced phenotype (Rosenbaum *et al*, 2011). Xport-B mutants also presented light enhanced retinal degeneration and extremely reduced protein levels of Rh1 and TRP. Interestingly, overexpression of Xport-A in the Xport-B mutant, or overexpression of Xport-B in the Xport-A mutant failed to rescue the ERG defects observed in these mutants, indicating that the roles of Xport-A and Xport-B are not redundant (Chen *et al*, 2015).

Rh1 also endures other post-translational modifications, such as chromophore binding. The chromophore of *Drosophila* Rh1 is made from β-carotene that is uptaken in the diet (Ozaki *et al*, 1993) and then processed to vitamin A, which is subsequently converted into 11-*cis* 3-hydroxy-retinal (Wang *et al*, 2007; Wang & Montell, 2007). The lack of chromophore incorporation in the *Drosophila* Rh1, by carotenoid dietary restriction, leads to extremely reduced protein levels of Rh1 (Ozaki *et al*, 1993). Furthermore, carotenoid deprivation affects Rh1 deglycosylation and transport, resulting in glycosylated Rh1 that appears to be retained in the ER presumably due to folding defects (Ozaki *et al*, 1993; Huber *et al*, 1994).

## Results and Discussion

### Rh1 TMD1-5 is double glycosylated in N20 and N196 in Xport-A^1^ homozygous mutants

Recently we have shown that the ER membrane protein complex (EMC) is required for the biogenesis and membrane insertion of Xport-A (Gaspar *et al*, 2022). In the last figure of that manuscript, we expressed 3 truncations of Rh1 (TMD1, TMD1-3 and TMD1-5) (Hiramatsu *et al*, 2019) in the background of flies heterozygous or homozygous for XportA^1^ mutation. Surprisingly, only Rh1 TMD1-5 was “affected” in the Xport-A^1^ homozygous flies, presenting itself as double glycosylated in N20 and N196, in contrast to single N20 glycosylation of Rh1 TMD1-5 in the Xport-A^1^ heterozygous background.

Rh1 contains two possible glycosylation sites: N20, within the N-terminal region and N196, in the second extracellular loop, between TMD4 and TMD5 (Fig. 1) (O’Tousa, 1992; Katanosaka *et al*, 1998; Webel *et al*, 2000). Although *in vitro* experiments with mammalian microsomes, have shown that Rh1 can be glycosylated at both sites (Katanosaka *et al*, 1998), only glycosylation at N20 has been shown to occur *in vivo* in WT and ninaA mutant flies (O’Tousa, 1992; Katanosaka *et al*, 1998; Webel *et al*, 2000). Nonetheless, mutation of the asparagine residue at both glycosylation sites (N20 and N196) to isoleucine (N20I and N196I) interferes with biogenesis of mutant and WT Rh1, resulting in the ER retention of Rh1 (Webel *et al*, 2000). Consequently, *Drosophila* eyes expressing Rh1 N20I or Rh1 N196I present large accumulations of Rh1 in the ER and Golgi membranes (Webel *et al*, 2000). Furthermore, as less Rh1 reaches the rhabdomeres, these mutant (Rh1 N20I or Rh1 N196I) eyes also present late onset age-related retinal degeneration (Webel *et al*, 2000).

**Figure 1.**
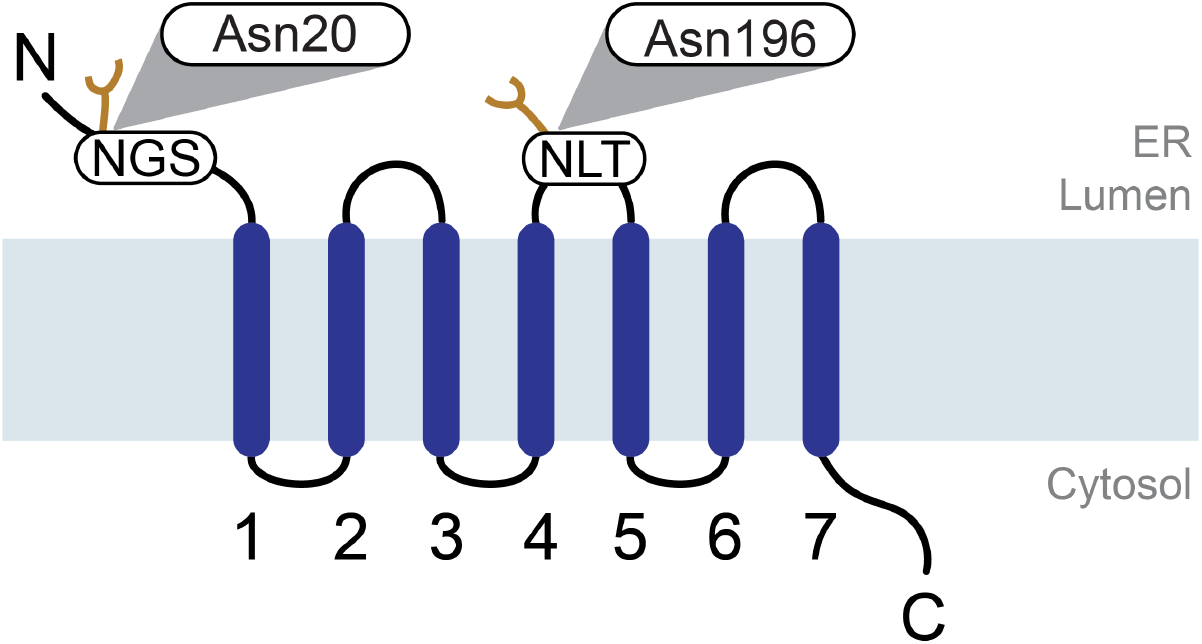
Schematic model of *Drosophila* Rhodopsin-1 showing the two N-glycosylation sites. Rhodopsin contains two predicted glycosylation sites, NGS (Asparagine-Glycine-Serine) in the N-terminal region, and NLT (Asparagine-Leucine-Threonine) in the second extracellular loop region, between TMD4 and TMD5. The predicted first glycosylation site is mapped to Asparagine at position 20 (N20) and the second glycosylation site is mapped to Asparagine at position 196 (N196). Sugar residues are represented in yellow.

While glycosylation of Rh1 in N20 seems to be necessary for maturation and/or transport in the secretory pathway (Webel *et al*, 2000), mature *Drosophila* Rh1 is completely deglycosylated. During maturation, Rh1 fully glycosylated 40kDa form is first trimmed in the ER to a 39kDa partly deglycosylated form and then is further de-glycosylated in the Golgi to a completely or almost completely de-glycosylated protein (Satoh *et al*, 1997; Rosenbaum *et al*, 2014). The higher molecular weigh of Rh1 in flies deprived of β-carotene seems to be due to glycosylation at N20 (Katanosaka *et al*, 1998).

### Xport-A TMD accommodates in Rh1 TMD1-5 but not in the full length Rh1 protein

Based on our findings in (Gaspar *et al*, 2022), we hypothesized whether Xport-A could be required for Rh1 biogenesis at the stage when the first 5 TMDs of Rh1 are inserted into ER membrane, rather than later, when all TMDs of the full length (FL) Rh1 are inserted into the membrane. To provide insights into this issue we resorted to AlphaFold2 (Jumper *et al*, 2021; Varadi *et al*, 2022) structural predictions of the Xport-A TMD together with full length (FL) Rh1 or Rh1 TMD1-5 (Fig. 2). AlphaFold2 computes five models and we display the topranked model (rank 1) for each complex in Fig 2. When bound to FL Rh1 (Fig 2A), the predicted structure of Xport-A TMD has low certainty standards (pLDDT values per residue) but an overall high pLDDT (>85) in complex with Rh1 TMD1-5 (Fig. 2B). Moreover, Xport-A/FL Rh1 structural predictions also display high *inter-complex predicted alignment errors* (PAE) and less favorable contacts, with 3 out of the 5 poses showing Xport-A TMD in a reverse topology (N terminal in the ER lumen and C terminal in the cytosol) from what is expected, since Xport-A is a TA protein (N terminal in the cytosol and C terminal in the ER lumen). Alphafold2 models are therefore consistent with Xport-A binding to Rh1 TMD1-5 rather than to FL Rh1. This is functionally supported by comparing the structure of FL Rh1 by itself (Fig.3A) with the pose of Xport-A TMD together with Rh1 TMD1-5 (Fig.3B); one can visualize that the Xport-A TMD partially overlaps with the would-be locations of TMD6, TMD7 and α-helix 8 of Rh1. Hence, Xport-A could function as a chaperone transiently, by mimicking the structural features of Rh1 TMD6, TMD7 and α-helix 8, when only the first 5 TMDs of Rh1 are inserted into the membrane. Moreover, Xport-A could be important for shielding Rh1 TMD1-5 Asp96, which otherwise would be unfavorably exposed to the membrane core.

**Figure 2.**
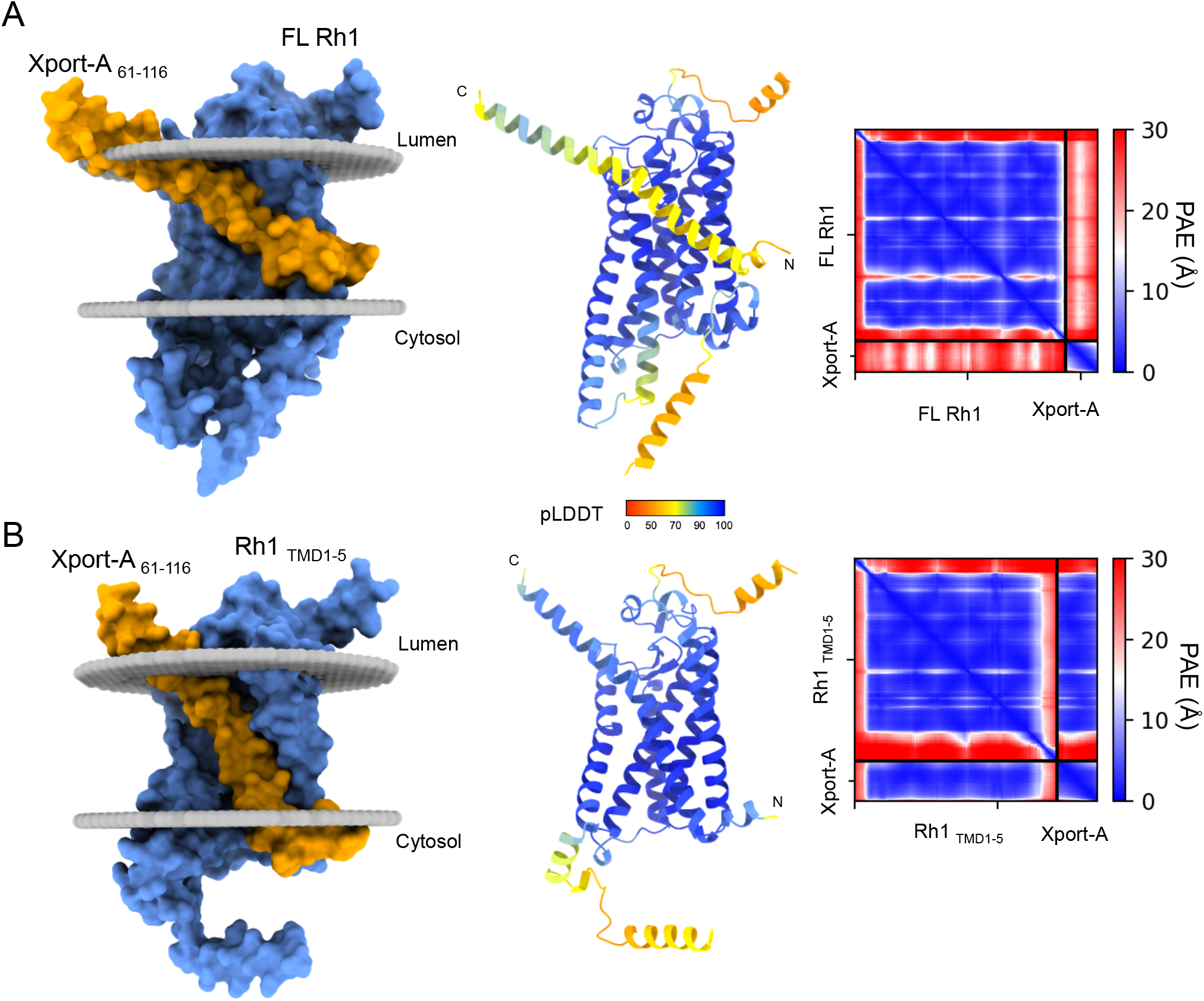
Xport-A TMD accommodates in Rh1 TMD1-5 but not in the full length Rh1 protein. AlphaFold2 structural predictions of the Xport-A together with (A) full length Rh1 or (B) Rh1 TMD1-5 (M1 to V241 +H333-A373). We display the top-ranked model for each complex, of the five models computed by AlphaFold2. To the left, Xport-A (amino acids 61 to 116) is represented in yellow and Rh1 FL and Rh1 TMD1-5 are in blue. In the middle, AlphaFold2 produces an estimate of confidence for each amino acid residue (pLDDT - Local Distance Difference Test), color-coded on a scale from 0 - 100. Values of pLDDT > 90 (blue) are expected to be modeled with high accuracy. To the right are represented the Predicted Aligned Errors (PAE) for each of the structure predictions.

**Figure 3.**
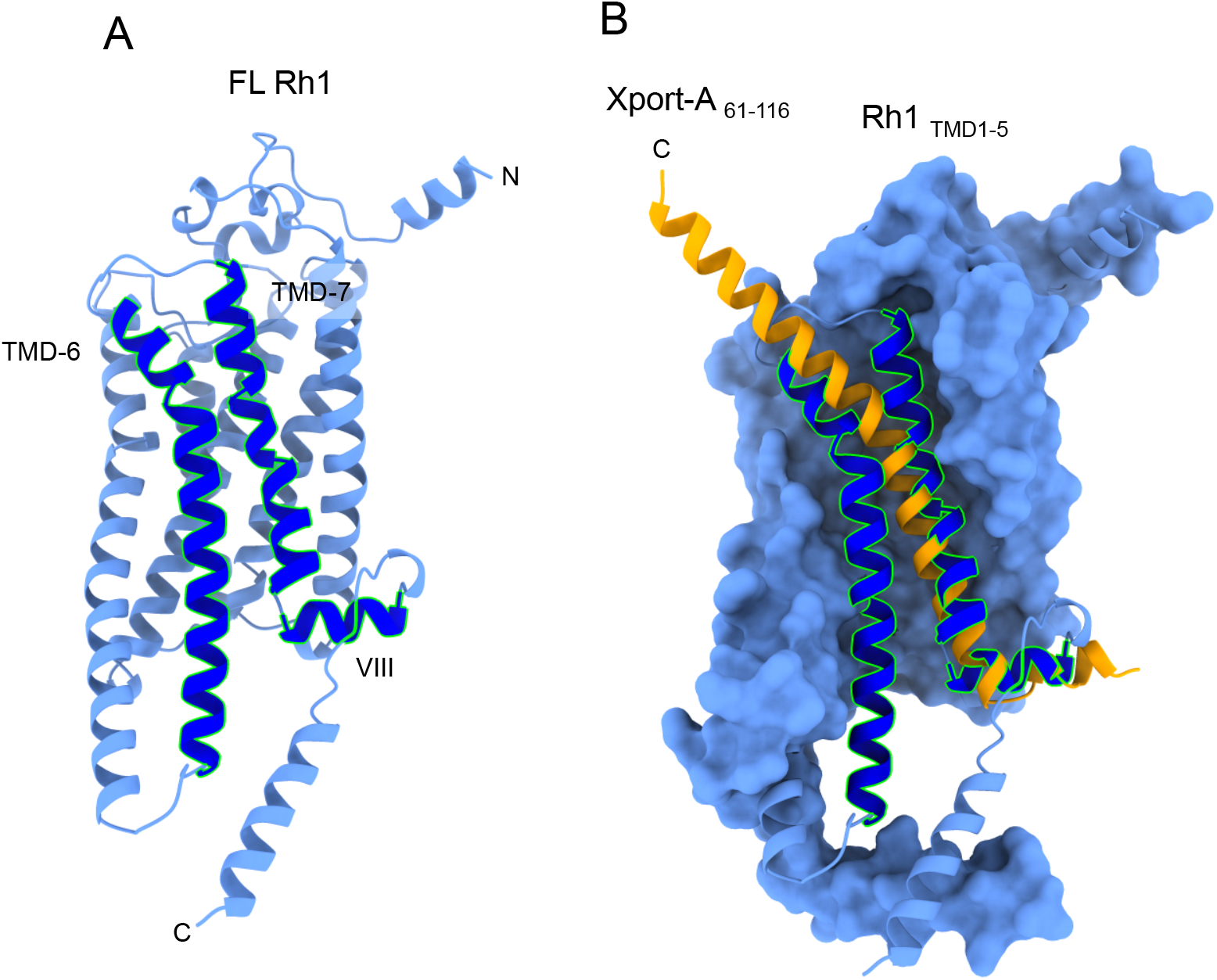
Xport-A overlaps with TMD6, TMD7 and α-helix 8 of Rh1. (A) Representation of FL Rh1 with TMD6, TMD7 and α-helix 8 highlighted in darker blue with green outlines. (B) Superimposition of Xport-A (amino acids 61 to 116 - in yellow) with TMD6, TMD7 and α-helix 8 of Rh1, with Rh1 TMD1-5 surface in pale blue.

### Prediction of structural interactions between Xport-A and Rh-1

Next we focus on some possible interactions between Xport-A and Rh1, based on the most likely pose that we described in Fig. 2B. Of particular interest, are the positions of the amino acid residues that we previously mutated to Leucine (Gaspar *et al*, 2022). Xport-A 1L, 2L, 3L and 4L (Fig. 4A) are a series of mutants which lead to an increasingly more hydrophobic TMD of Xport-A, progressively bypassing the EMC requirement for membrane insertion. We have also shown that Xport-A 2L and 4L rescue the expression of Rh1 in EMC mutant cells (Gaspar *et al*, 2022), but in both cases these rescues were only partial, suggesting that these 2L and 4L mutants have reduced function, although they are inserted into the membrane, even in EMC mutant cells (Gaspar *et al*, 2022). In Fig. 4B, we show the position of the 4 amino acid residues (N83, T84, T90 and H95) that we mutated to leucine in Xport-A 4L and all of them are oriented into the interface, in a position to interact with amino acid residues in Rh1 TMD1-5. In fact, we could observe that Xport-A 2L is worse than Xport-A (WT) but better than Xport-A 4L at rescuing the expression of Rh1 (and TRP) in fly eyes homozygous for the Xport-A^1^ mutation (Fig 5). This result suggests that the biological functionality of Xport-A 2L is less compromised than in Xport-A4L. This again supports a role of Xport-A as a polarity shield to Rh1 TMD1-5, which is compromised when its own TMD’s polar residues (N83, T84 and T90) become mutated to apolar ones.

**Figure 4.**
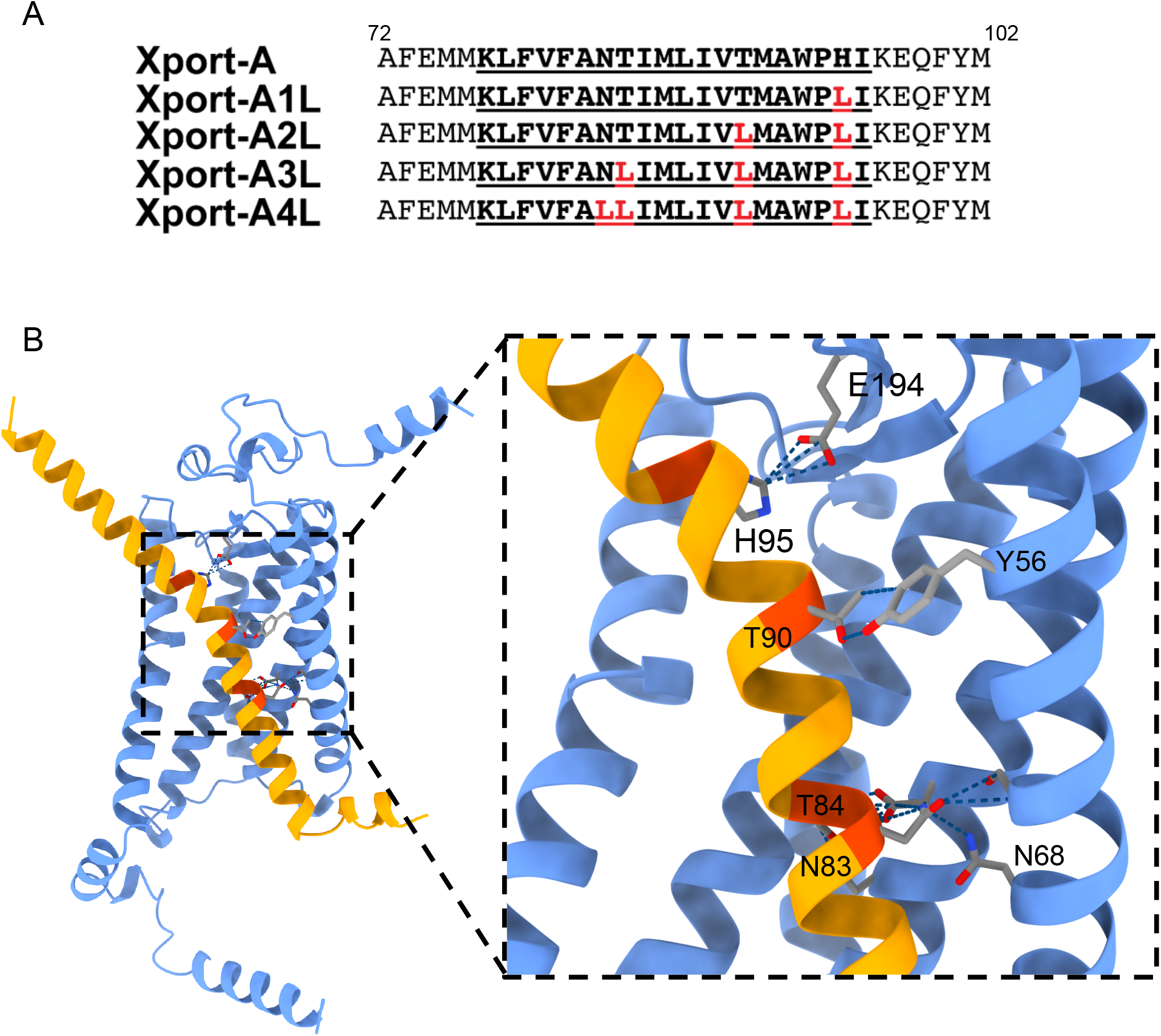
Interactions between Xport-A TMD and Rh-1 TMD1-5. (A) Amino acid sequence alignments of the TMDs (bold, underlined) of Xport-A, Xport-A1L, Xport-A2L, Xport-A3L and Xport-A4L. (B) Prediction of interactions between the Xport-A amino acid residues that were mutagenized to L (N83, T84, T90 and H95) and Rh1.

**Figure 5.**
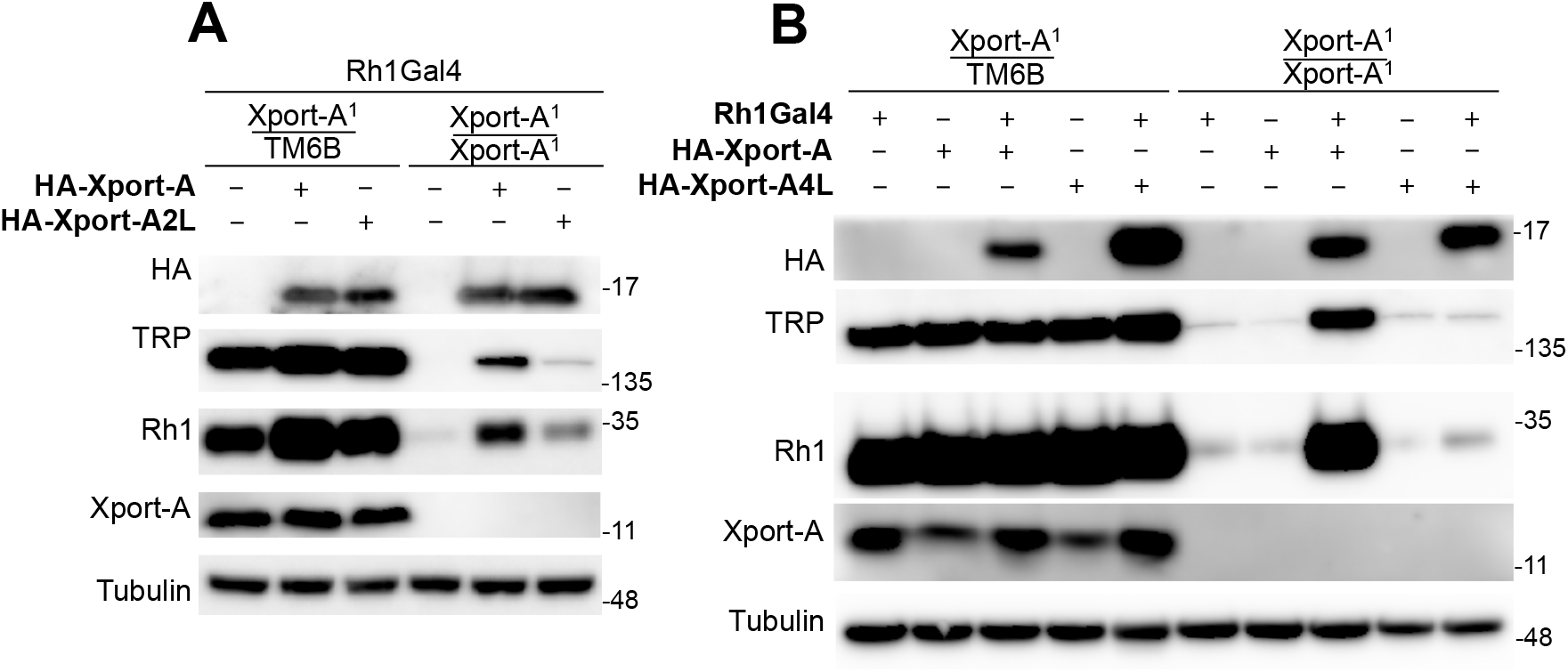
Rescue of Rh1 and TRP in Xport-A1 homozygous eyes by Xport-A, Xport-A2L and Xport-A4L. (A) Immunoblot of fly heads expressing HA-Xport-A or HA-Xport-A2L in Xport-A heterozygous (Xport-A^1^ /TM6B) or homozygous mutant flies (Xport-A^1^/Xport-A^1^), (B) Immunoblot of fly heads expressing HA-Xport-A or HA-Xport-A4L in Xport-A heterozygous (Xport-A^1^ /TM6B) or homozygous mutant flies (Xport-A^1^ /Xport-A^1^). In both (A) and (B) the blots were probed with antibodies against HA, TRP, Rh1, Xport-A and Tubulin, the UAS constructs were expressed under the control of Rh1-GAL4 and each lane was loaded with protein extracts from approximately 2,7 fly heads.

Furthermore, there are additional interactions between amino acid residues in Rh1 and the C-terminal domain of Xport-A, that projects into the lumen of the ER (Fig. 6). For example, H95 of Xport-A interacts with E194 of Rh1 (Fig. 6), which could be important in keeping the beta-loop-beta (Rh1 Y191 to I202) motif deep in the plane of the membrane and protected by the N terminal “crown” of 3 small α-helices (Rh1 S22 to Q41), which also interacts with amino acids in the ER luminal C-terminal domain of Xport-A (Fig. 6 inset); Xport-A Y101 and Q105 interact with Rh1 F42 and Q41, respectively. Of note, N20 is well exposed to ER luminal glycosylating enzymes, while N196 is not, since it is within the protected beta-loop-beta motif. So, in order for N196 to be accessible for glycosylation (in Xport-A mutants), some dramatic misfolding of the N terminal “crown” and/or beta-loop-beta motif must occur. The chaperone role of Xport-A might therefore extend to the protection of the beta-loop-beta motif. Finally, we would like to highlight that the structure of the beta-loop-beta motif must be important for Rh1 biogenesis, since at least 4 mutations in this motif (E194Q, E194K, G195S, C200Y) have been reported to cause reduced biogenesis of Rh1 and retinal degeneration (Colley *et al*, 1995; Kurada & O’Tousa, 1995; Zheng *et al*, 2015).

**Figure 6.**
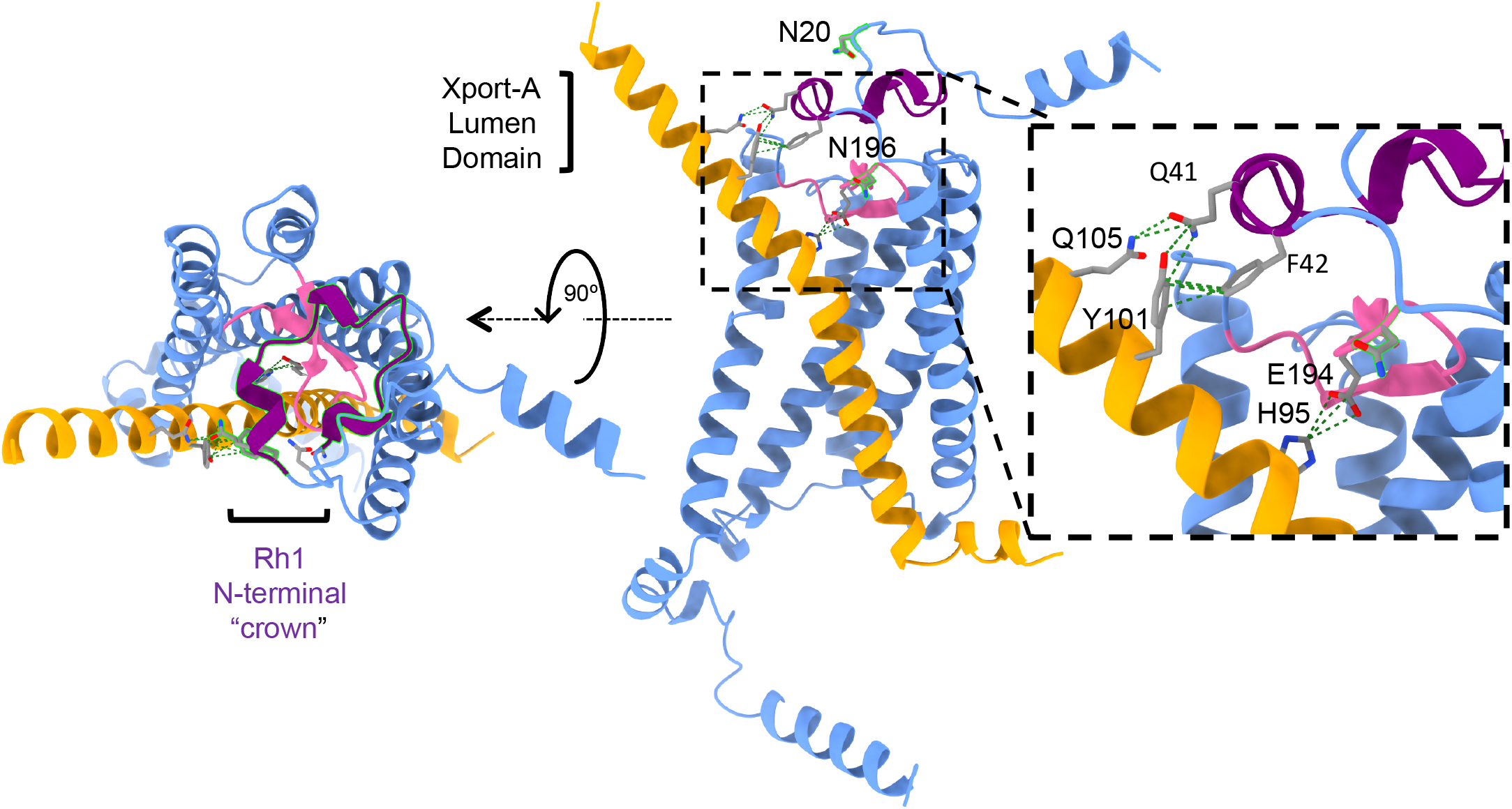
Interactions between Xport-A ER lumen domain and Rh-1 TMD1-5. Xport-A (amino acids 61 to 116) is in yellow. Rh1 TMD1-5 is in pale blue. The N-terminal “crown” of Rh1 (S22 to Q41) is in purple. The beta-loop-beta of Rh1 (Y191 to I202) is in pink.

### Molecular dynamics simulation of the interaction between Rh1 and Xport-A

In order to provide an independent confirmation of the interactions predicted above with AlphaFold2, we performed molecular dynamics (MD) simulations, using the Martini 3 coarse-grained (CG) model (Souza *et al*, 2021). Rh1 TMD1-5 and Xport-A were simulated in an ER membrane mimic, and left to interact spontaneously. As shown in Fig. 7A, all three independently performed replicates converged to an Xport-A/Rh1 TMD1-5 interaction analogous to the pose predicted by AlphaFold2 (Fig. 2B). In this interaction Xport-A again serves as chaperone for Rh1 TMD1-5, shielding the exposed polar/charged residues of Rh1 TMD1-5. In contrast, when initially-bound simulations were done with Rh1 TMD1-5 and Xport-A4L (Fig. 7B) an obviously weaker binding was observed, although Xport-A4L never dissociated completely from Rh1 TMD1-5. Supporting the stability of the Xport-A/Rh1 TMD1-5 interaction, simulations started with both proteins bound in the AlphaFold2-predicted pose remained tightly bound for the entirety of the multi-μs simulated timescale (Fig. 7C).

**Figure 7.**
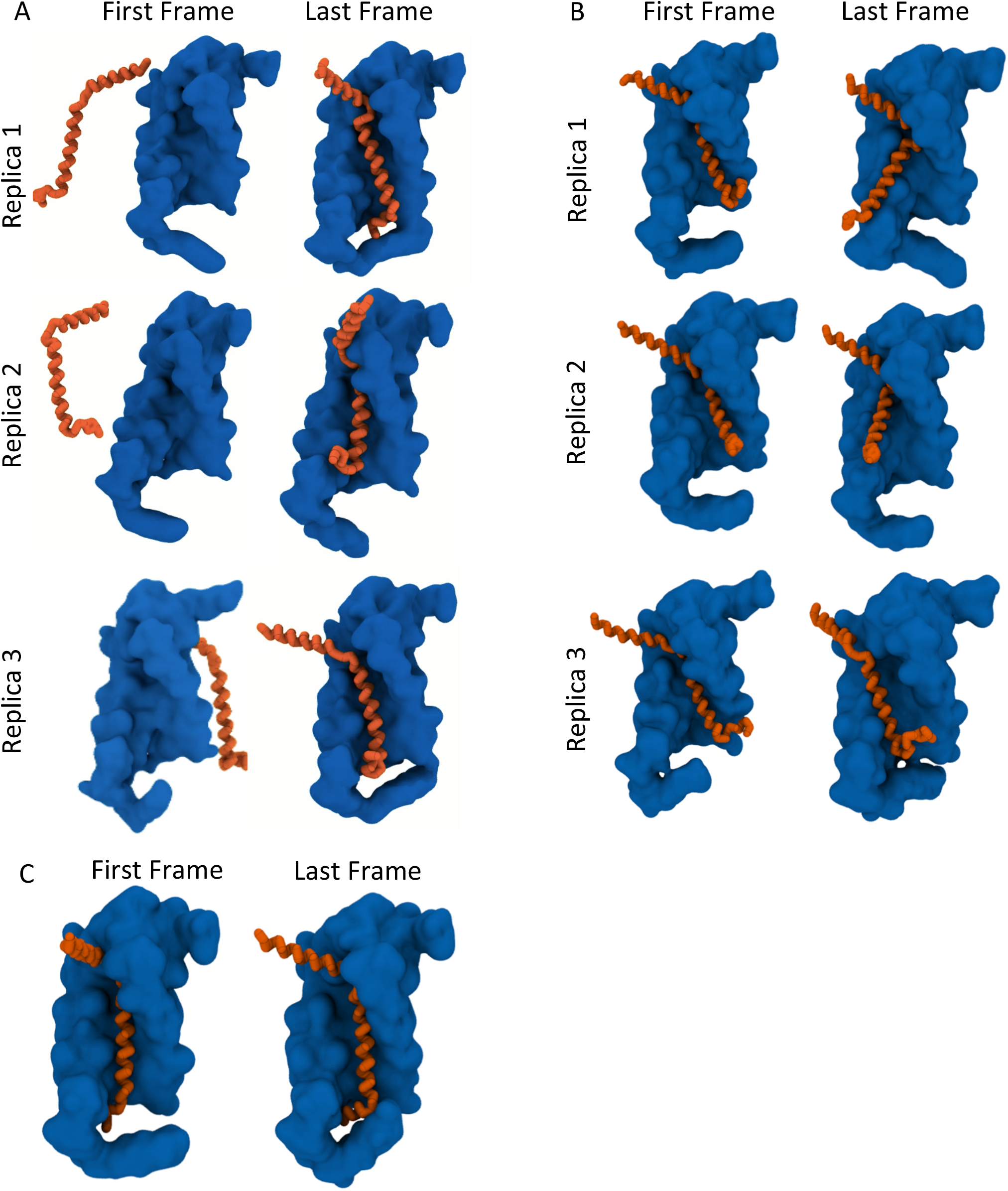
CG MD simulations of the protein-protein interactions between Rh1 TMD 1-5 and Xport-A. Simulations were done using the Martini 3 CG model with Rh1 TMD 1-5 (surface representation in blue) and Xport-A (backbone stick representation in orange). In the left panels are the first frames of the simulations and on the right the last frames. All replicates were run for at least 13μs (A) Simulations set up with Rh1 TMD 1-5 and Xport-A inserted separately in the membrane. (B) Simulations with Rh1 TMD1-5 and Xport-A 4L inserted in the membrane as a complex. (C) Simulations with Rh1 TMD1-5 and Xport-A inserted in the membrane as a complex; only one of the replicates is shown, with a representative bound behavior for the entire simulation.

### Conclusions

Based on the results of (Gaspar *et al*, 2022) and the results described here we favor a model where Xport-A acts as chaperone during the biogenesis of Rh1, at a transient step, when the TMDs 1-5 of Rh1 are already inserted in the ER membrane, but TMDs 6-7 are not yet inserted or at least not yet present in Rh1 structure. Interactions of Xport-A with Rh1 TMD1-5 must be essential to stabilize the TMDs of Rh1 but also to stabilize the N terminal ER luminal domain of Rh1 and the beta-loop-beta motif between TMD4 and TMD5, allowing for the correct folding and biogenesis of Rh1.

## Methods

### *Drosophila* stocks

Flies and crosses were raised with standard cornmeal fly food, at 25°C under 12 h light/12 h dark cycles. HA-Xport-A, HA-Xport-A2L and HA-Xport-A4L were described in (Gaspar *et al*, 2022). Xport-A^1^ mutation was described in (Rosenbaum *et al*, 2006).

### AlphaFold modelling of Xport-A Rh1 complex

We used AlphaFold-Multimer (Evans *et al*, 2022) to predict binding interfaces, a refined version of AlphaFold2 (Jumper *et al*, 2021) for complex prediction. As a first stage, we used the sequences of Rh1 and Xport-A TMDs as input to predict the 3D structure of Xport-A bound to Rh1. Next, we run independent predictions replacing the full length Rh1 with the sequence of Rh1 TMD1-TMD5 construct, including the C-Terminus of Rh1 (M1 to V241 + H333-A373) and a V5 tag. We did not use template structures for the predictions iterated for up to 48 recycles, followed by energy refinement with AMBER using default settings implemented in LocalColabFold (Mirdita *et al*, 2022) and using MMseqs2 for creating multiple sequences alignments (Steinegger *et al*, 2017). Model confidence was assessed by the predicted Local Distance Difference Test (pLDDT) and inter-complex predicted alignment error (PAE), i.e., the uncertainty about the interface. Regions with pLDDT > 90 are expected to have high accuracy.

### Immunoblotting of fly heads

Heads from 1-day old flies were homogenized in 2xLDS buffer +DTT with a pellet pestle, and then diluted with MilliQ water. Protein denaturation was performed by incubating extracts at 65 C (15 min). Samples were run in SDS-PAGE, transferred to PVDF membranes (Amersham Hybond) and probed with the following antibodies: mouse anti-V5 (1:1,000) (R960-25, Invitrogen), mouse anti-TRP (1:300) (Mab83F6, DSHB), mouse anti-Rh1 (1:200) (4C5, DSHB), rat anti-Xport-A antibody (1:400) (kind gift of Craig Montell), and mouse anti-tubulin (1:1,000) (AA4.3, DSHB).

### Molecular dynamics simulation of the interaction between Rh1 and XportA

The membranes were built as described in (Wassenaar *et al*, 2015), adapting the lipid composition to what has been described for the membrane of the endoplasmic reticulum described in CHARMM-GUI (Jo *et al*, 2015). The Coarse Grain structure of the Rh1 TMD1-5, Xport-A and Xport-A4L proteins was built using the Martini method as described (Souza *et al*, 2021). The simulations were done by adding Rh1 TMD1-5 and Xport-A separately to the membrane, or adding a pre-made Rh1 TMD1-5/Xport-A complex to the membrane or adding the Rh1 TMD1-5/Xport-A4L complex to the membrane. Three separate replicates were generated by doing separate equilibration steps.

### Molecular dynamics simulation of the interaction between Rh1 and XportA

The CG topology and structure of the Rh1 TMD1-5, Xport-A and Xport-A 4L proteins was built using the *martinize2* script (https://github.com/marrink-lab/vermouth-martinize), employing elastic network restraints for maintaining the Rh1 TMD1-5 structures, but only main-chain angle/torsion potentials in maintaining Xport-A’s structure. Xport-A’s secondary structure was assumed to be entirely α-helical with the exception of termini, proline residues, and proline-flanking residues. Membranes were built and proteins inserted using the *insane* script (Wassenaar *et al*, 2015), adapting the lipid composition to that of the endoplasmic reticulum membrane (Jo *et al*, 2015). Rh1 TMD1-5 and Xport-A were either inserted separately in the membrane, or as a pre-made Rh1 TMD1-5/Xport-A complex. For each system, three separate replicates were performed of at least 13μs each. We used the GROMACS simulation package version 2020 (Abraham et al, 2015). Lennard-Jones interactions were cutoff at 1.1 nm; Coulombic interactions were treated, with the same cut-off, using reaction-field electrostatics with a dielectric constant of 15 and an infinite reaction-field dielectric constant. Temperature was kept at 300 K by a v-rescale thermostat with a coupling time of 4.0 ps. Pressure was coupled semi-isotropically at 1.0 bar to a Parrinello-Rahman barostat, with a relaxation time of 16.0 ps. Simulations were run at a 20 fs time step. Visualization and rendering of the simulations were performed with the molecular graphics viewer VMD version 1.9.3 (Humphrey *et al*, 1996).

## Acknowledgements

The authors acknowledge Fundação para a Ciência e a Tecnologia, I.P. (FCT) for funding Project MOSTMICRO-ITQB, with references UIDB/04612/2020 and UIDP/04612/2020. MNM further acknowledges FCT for fellowship CEECIND/04124/2017 and TNC is the recipient of the grant CEECIND/01443/2017. The project leading to these results was funded by the grants LCF/PR/HR17/ 52150018 (‘la Caixa’ Foundation) and FCT AGA-KHAN/541141368/2019 (FCT and Aga Khan Foundation).

## Notes

### Competing Interest Statement

The authors have declared no competing interest.

